# ProtLoc-GRPO: Cell line-specific subcellular localization prediction using a graph-based model and reinforcement learning

**DOI:** 10.1101/2025.07.17.665451

**Authors:** Shuai Zeng, Weinan Zhang, Chaohan Li, Yuexu Jiang, Duolin Wang, Qing Shao, Dong Xu

**Author notes:** Email: Shuai Zeng, Weinan Zhang, Chaohan Li, Yuexu Jiang, Duolin Wang, Qing Shao, Dong Xu.

## Abstract

Subcellular localization prediction is crucial for understanding protein functions and cellular processes. Subcellular localization is dependent on tissue and cell lines derived from different cell types. Predicting cell line-specific subcellular localization using the information of protein-protein interactions (PPIs) offers deeper insights into dynamic cellular organization and molecular mechanisms. However, many existing PPI networks contain systematic errors that limit prediction accuracy. In this study, we propose a reinforcement learning approach, ProtLoc-GRPO, to enhance subcellular localization prediction by optimizing the structure of the underlying PPI network. ProtLoc-GRPO learns to rank and retain the most informative PPI edges to maximize the macro-F1 score for cell line-specific subcellular localization. Our approach yields a 7% improvement in macro-F1 score over the baseline. We further evaluate its robustness across various edge pruning rates and benchmark it against conventional pruning strategies. Results show that our proposed method consistently outperforms existing approaches. To our knowledge, this work represents the first study to predict cell line-specific protein subcellular localization and the first application of the Group Relative Policy Optimization (GRPO) framework to a graph-based model for bioinformatics tasks.

## Introduction

Protein function and regulation are highly dependent on subcellular localization. Eukaryotic cells are compartmentalized into various organelles, each providing distinct chemical environments and specific sets of interaction partners essential for proper protein activity^1^. This intricate organization ensures that proteins are precisely positioned where they can perform their designated roles. Accurate localization ensures proteins engage in appropriate interactions and participate effectively in cellular processes such as signaling and metabolism, orchestrating complex cascades and maintaining cellular homeostasis^2^. Conversely, even subtle mislocalization can disrupt these finely tuned processes, leading to protein dysfunction and is often associated with various diseases, including neurodegenerative disorders^3^, metabolic syndromes^4^, and cancer^5,6^. Protein subcellular localization is a complex process regulated by multiple factors, such as signal peptides, protein-protein interactions (PPIs), protein folding, and alternative splicing^7^.

Proteins can exhibit different localizations across different cell types^8,9^. This variability makes predicting and understanding protein subcellular localization particularly challenging. Over the past decades, the Human Protein Atlas (HPA)^1,10,11^ has dedicated significant effort to mapping the spatial distribution of human proteins across various cells, tissues, and organs. As of its latest release, the HPA has annotated the subcellular localization of proteins encoded by over 13,000 genes, covering approximately 65% of the human protein-coding genome. However, each protein has been profiled in up to only three of the 37 available cell lines using immunofluorescence, creating a substantial gap in our understanding of protein localization across diverse cellular contexts. This limitation hinders large-scale, cell line-specific insights into protein subcellular localization, function and dynamics.

Numerous advanced deep learning models have been developed to predict subcellular localization with high accuracy, relying on identifying motifs and patterns within protein sequences to infer their subcellular locations, such as DeepLoc^12^ and MULocDeep^13^. Other notable approaches have leveraged transfer learning by fine-tuning large protein language models for protein localization, such as ProtGPS^14^, PEFT-SP^15,16^, TargetP^17^, and the SignalP^18–20^ series. Recent works^21,22^ have improved subcellular localization prediction by combining features from PPIs, GO annotations^23,24^, and protein sequences. However, all these available computational methods primarily focus on bulk-level localization rather than cell line-specific localization, limiting their ability to accurately identify essential proteins across diverse cell lines derived from different cell types.

Protein interactions are closely related to the localization of proteins^25,26^. In other words, for proteins to interact, they necessarily share a common subcellular localization or, at least transiently or conditionally, an interface between two physically adjacent compartments. However, PPI data inherently contain limitations and noise. Some interactions^27,28^ are detected only under specific experimental conditions or *in vitro* environments and may not accurately reflect interactions that occur *in vivo*. A notable example is the Yeast Two-Hybrid (Y2H) screening^29,30^, a widely used technique for detecting binary PPIs. While Y2H is high-throughput and useful for identifying novel interactions, it operates in the nucleus of yeast cells and may force proteins into non-native compartments or conformations, potentially producing interactions that would not occur in physiological settings^30,31^. Moreover, interactions discovered through Y2H often lack post-translational modifications or co-factors present in native environments, further limiting biological relevance. Additionally, variability across experimental platforms and technical artifacts from high-throughput screening methods can lead to false positives or non-physiological interactions^32^. These inaccuracies can cause misleading outcomes in downstream analyses, including protein subcellular localization prediction.

To address this issue, we propose leveraging machine learning techniques to filter PPI edges, effectively distinguishing interactions that contribute positively to localization prediction from those that may introduce noise or errors. By training models to identify and retain only the most informative PPIs, we can enhance both the accuracy and robustness of protein subcellular localization predictions. Precisely, we frame our work as pruning low-importance PPI edges to improve protein subcellular localization using reinforcement learning (RL) and graph-based model. Thus, we propose an RL-based PPI edge pruning method, ProtLoc-GRPO, using the Group Relative Policy Optimization (GRPO)^33^ framework. GRPO has been widely adopted for optimizing large language models (LLMs). To the best of our knowledge, our work is the first application of GRPO in the bioinformatics domain.

ProtLoc-GRPO consists of two components: a Graph Attention Network (GAT)^34^ policy model that ranks the importance of PPI edges for subcellular localization prediction, and a model-based reward function that provides feedback to guide the RL policy model toward optimal pruning decisions. The policy model prunes low-importance edges according to a predefined pruning rate, retaining only the most informative ones. These pruned PPI networks are then used to construct cell line-specific graph-based models for protein subcellular localization. Experimental results show that our method improves macro-F1 score by up to 7% compared to models trained on unpruned PPI networks. We further evaluate ProtLoc-GRPO across multiple pruning rates and using different graph-based backbones. In all scenarios, our method consistently outperforms the non-pruned baselines that use all PPI edges to build graph-based classification models, demonstrating its effectiveness in identifying critical PPI edges for enhancing subcellular localization prediction. Importantly, our method approach is not only valuable within the context of our study but also broadly applicable to other bioinformatics challenges, such as protein function annotation and the investigation of disease mechanisms.

## Results

### Constructing a protein interaction network

We assembled a dataset containing protein sequences and cell line-specific subcellular localization annotations from HPA. To ensure data quality and statistical robustness, we first analyzed the distribution of proteins with localization annotations across all 37 available cell lines (see Supplementary Figure 1). Our analysis revealed that most cell lines have a limited number of annotated proteins, making them unsuitable for reliable modeling. Therefore, we selected only those cell lines with more than 1,000 proteins annotated with subcellular localization information, yielding a final set of 5 cell lines. These selected cell lines are derived from diverse tissue origins, such as lung, kidney, and skin (see Supplementary Table 1). After identifying eligible proteins in these cell lines, we mapped them to a unified PPI network, which was constructed by integrating curated interactions from BioGRID^27,35^, HuRI^36^, STRING^28^, and the Menche et al. datasets^37^ (Figure 1a). We extracted the subgraph induced by eligible proteins and retained the largest connected component, resulting in a globally unified PPI, which serves as the basis for subsequent training, pruning, and localization prediction tasks.

**Table 1.** Comparison of graph-based models constructed using unpruned PPI networks versus ProtLoc-GRPO pruned PPI networks. The reported metrics are presented as percentages, along with their standard deviations. Bold values indicate the best performance for each metric.

| Method | Prune rate | Reward function/<br>test model | Mean AUC | Macro-F1 | Macro-precision | Macro-recall |
| --- | --- | --- | --- | --- | --- | --- |
| Unpruned | 0% | GAT | 71.06 ± 1.02 | 35.00 ± 1.24 | 31.16 ± 2.32 | 56.22 ± 4.56 |
| | | GCN | 71.46 ± 0.49 | 35.46 ± 0.73 | 33.24 ± 0.89 | 54.32 ± 3.34 |
| | | GraphSAGE | 71.20 ± 0.75 | 35.22 ± 0.65 | 31.30 ± 0.84 | 55.96 ± 2.33 |
| ProtLoc-GRPO | 10% | GAT | 71.4 ± 0.16 | 37.22 ± 0.89 | 34.43 ± 4.34 | 56.40 ± 0.36 |
| | | GCN | 71.80 ± 0.39 | 37.45 ± 0.68 | 34.84 ± 2.31 | 56.66 ± 1.45 |
| | | GraphSAGE | **71.95 ± 0.30** | 37.59 ± 1.04 | 33.24 ± 0.67 | 55.93 ± 2.47 |
| | 15% | GAT | 71.68 ± 1.03 | 37.77 ± 0.88 | 35.22 ± 1.60 | 57.56 ± 1.27 |
| | | GCN | 71.69 ± 0.55 | 38.03 ± 0.91 | 35.57 ± 2.37 | 56.05 ± 2.39 |
| | | GraphSAGE | 71.73 ± 0.97 | 37.92 ± 1.12 | 35.10 ± 1.34 | 57.37 ± 3.41 |
| | 20% | GAT | 71.42 ± 0.85 | 37.16 ± 1.00 | 33.72 ± 1.43 | **61.82 ± 1.30** |
| | | GCN | 71.65 ± 0.92 | **38.11 ± 0.74** | 35.55 ± 2.15 | 56.95 ± 1.61 |
| | | GraphSAGE | 70.63 ± 1.04 | 36.67 ± 1.02 | 33.51 ± 1.36 | 57.90 ± 2.48 |
| | 25% | GAT | 71.21 ± 0.98 | 37.82 ± 1.44 | 34.83 ± 3.19 | 58.37 ± 1.79 |
| | | GCN | 70.62 ± 0.54 | 37.59 ± 0.24 | 33.53 ± 0.92 | 58.17 ± 0.60 |
| | | GraphSAGE | 71.43 ± 0.65 | 37.92 ± 0.96 | **35.92 ± 0.91** | 55.19 ± 4.92 |
| | 30% | GAT | 71.55 ± 0.60 | 37.69 ± 0.08 | 34.84 ± 2.12 | 56.39 ± 2.39 |
| | | GCN | 70.98 ± 1.42 | 37.48 ± 0.95 | 34.86 ± 1.73 | 58.40 ± 1.55 |
| | | GraphSAGE | 71.35 ± 1.02 | 37.69 ± 0.58 | 35.53 ± 2.64 | 56.54 ± 2.07 |
| | 40% | GAT | 70.88 ± 0.99 | 36.35 ± 1.62 | 32.54 ± 2.93 | 59.72 ± 0.74 |
| | | GCN | 70.44 ± 0.66 | 37.70 ± 0.51 | 34.78 ± 0.27 | 57.46 ± 0.56 |
| | | GraphSAGE | 70.51 ± 1.07 | 36.54 ± 0.75 | 32.35 ± 1.41 | 59.16 ± 2.40 |
| | 60% | GAT | 70.59 ± 1.42 | 36.17 ± 1.16 | 33.10 ± 2.67 | 60.33 ± 3.73 |
| | | GCN | 70.71 ± 0.70 | 37.38 ± 0.96 | 33.15 ± 2.22 | 60.67 ± 1.89 |
| | | GraphSAGE | 70.65 ± 0.84 | 37.29 ± 0.89 | 33.49 ± 1.11 | 60.95 ± 1.82 |
| | 80% | GAT | 69.95 ± 1.07 | 36.21 ± 1.59 | 32.59 ± 4.27 | 59.64 ± 2.36 |
| | | GCN | 69.75 ± 0.96 | 36.65 ± 0.69 | 33.02 ± 0.96 | 58.46 ± 2.78 |
| | | GraphSAGE | 70.16 ± 1.36 | 37.16 ± 0.95 | 33.47 ± 1.60 | 58.96 ± 2.38 |

**Figure 1.**
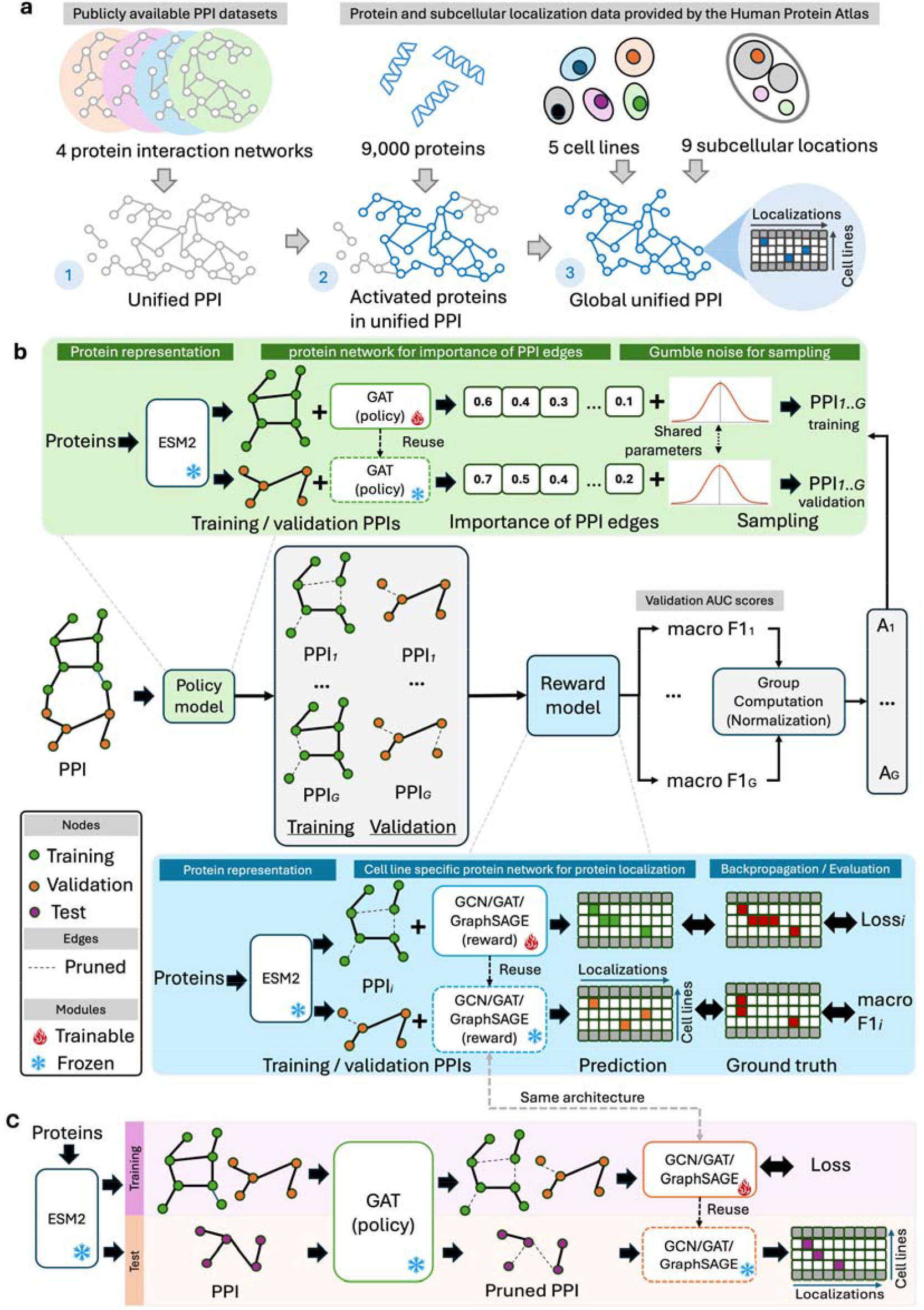
Overview of the subcellular dataset and ProtLoc-GRPO. (a) Globally unified PPIs are constructed from multiple different protein interaction networks. (b) The training process of ProtLoc-GRPO involves a GAT policy model that prunes low-importance PPI edges optimized by a reward function that returns the macro-F1 on the validation set. (c) During the test stage, the optimized GAT policy model deterministically prunes PPI edges without stochastic sampling. A graph-based model with the same architecture as that used in the reward function is then constructed and trained on the pruned PPI network from the training set, followed by final evaluation on the test set.

### Overview of ProtLoc-GRPO

Unlike conventional GRPO implementations, where the policy model is typically an LLM and rewards are computed through rule-based verification, the proposed ProtLoc-GRPO framework incorporates a GAT-based policy model and a task-specific reward function, and omits the KL divergence penalty term, as illustrated in Figure 1b. The GAT policy aims to identify the most important PPI edges and prune those with lower importance. The reward function consists of training and predicting with a graph-based model for subcellular localization, followed by a macro-F1 scoring function that compares the predicted localizations with the ground truth, providing reward signals to update the GAT policy model. The graph-based model in the reward function can be implemented using GCN^38^, GAT, or GraphSAGE^39^. For each training iteration, we used the ESM2 650M model to generate rich and informative protein representations as node features for both the GAT policy model and the graph-based model in the reward function. The GAT policy was built using a globally unified PPIs, while the graph-based model in the reward function is built based on the pruned PPI edges from the GAT policy model. We performed 5-fold cross-validation to evaluate the ProtLoc-GRPO, where the training and validation sets are used during reinforcement learning training, and the test set is used for final evaluation. Because the reward function involves both training and prediction stages, it requires access to both the training and validation sets. The architecture of GAT policy model and graph-based model in reward function are detailed in Figure 2.

**Figure 2.**
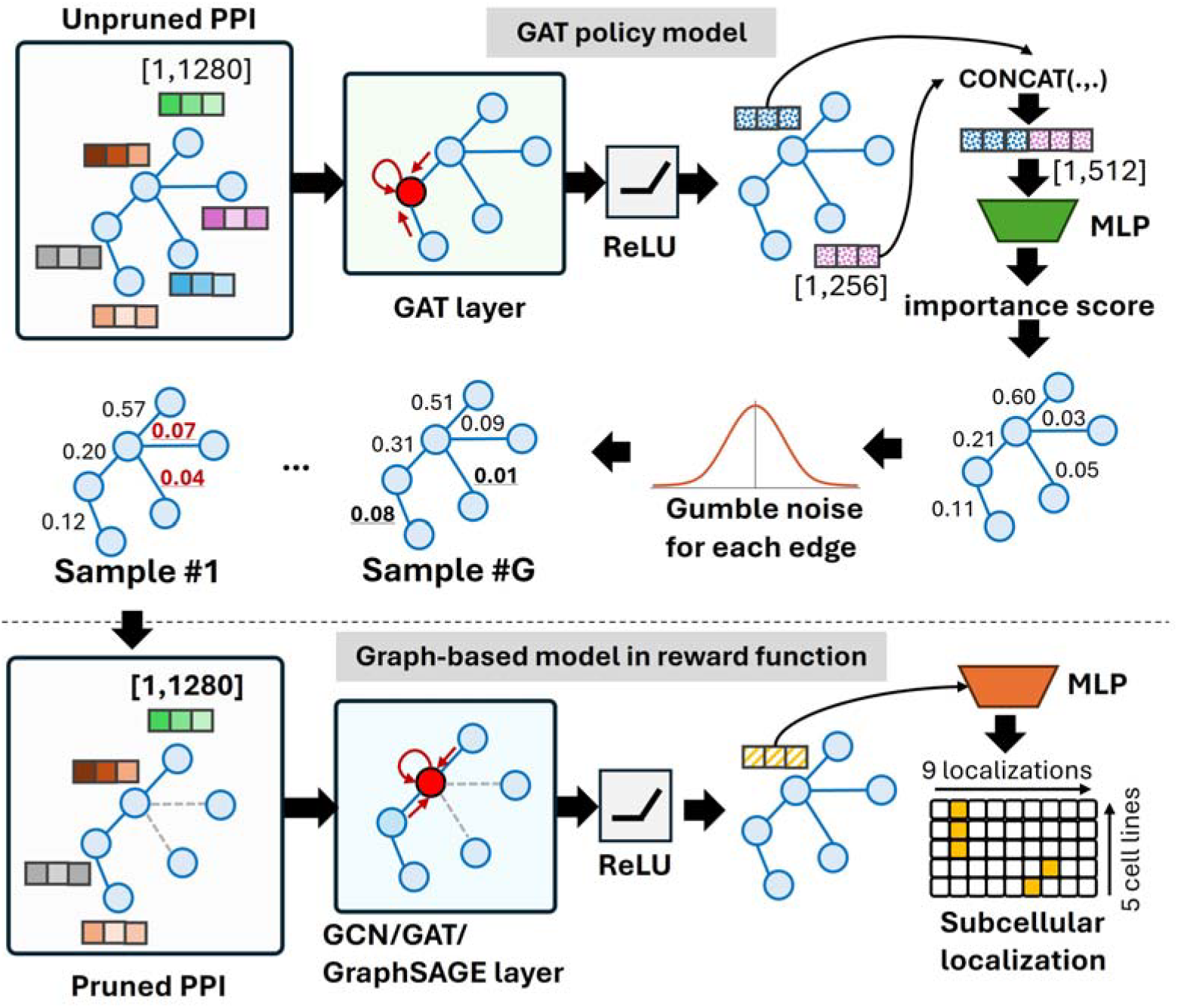
Architectures of the GAT policy model and the graph-based model in the reward function. All graph models take as input the mean-pooled protein representations generated by the ESM2-650M model. The GAT policy model uses concatenated embeddings of protein node pairs to compute importance scores for each PPI edge. In contrast, the graph-based model in the reward function performs node classification across five cell lines and nine subcellular localization categories.

In each optimization iteration, the GAT policy model estimates the importance of each PPI edge from the full edge set for protein subcellular localization prediction. Specifically, we treated each undirected PPI edge as two directed edges in opposite directions and computed the importance score for each direction separately. To generate *G* sampled PPI edge sets for the training and validation sets, Gumbel noise is added to the importance scores, and low-importance edges are pruned based on a predefined pruning rate. For each of the *G* pruned edge sets, a graph-based model is constructed and trained independently on the same training data. The macro-F1 score on the validation set is then used as the reward signal to guide the policy model toward identifying informative edges that generalize to unseen data, following strategies adopted in prior work^40^. The reward scores are normalized using the *z*-score normalization, resulting in positive advantage scores for high macro-F1 values and negative scores for low macro-F1 values. It is important to note that some proteins are annotated with subcellular locations in only up to three cell lines. Additionally, certain subcellular locations may have no corresponding proteins in the validation set. In such cases, we excluded those localizations from the macro-F1 calculation.

We borrowed the idea from the post-training step of LLM, in which the model is finetuned using a cold start dataset rather than starting from the base model, in order to reduce the instability of training and achieve better convergence^41^. We initialized the GAT policy model by training it on a cold start dataset, where attention values from a well-trained GAT on a unified PPI network served as supervision targets. During the RL training, the ProtLoc-GRPO optimizes the whole weights of the GAT policy model to adjust its edge importance estimates based on the reward signal from the reward function. The objective function of the ProtLoc-GRPO is to maximize the importance of edges that improve the macro-F1 score. At the inference stage, the policy operates in deterministic mode, selecting only the most important edges without stochastic sampling on the test set. These pruned PPI edges are then used to build a new randomly initialized graph-based model for subcellular localization, which is used for final evaluation. To ensure consistent performance, the graph-based model built on the pruned PPI edges adopts the same architecture as the graph-based model in the reward function. For example, if GCN is used in the reward function in ProtLoc-GRPO, the corresponding graph-based subcellular localization model in the test stage will also use GCN.

### Pruned PPI edges outperform non-pruned baselines for subcellular localization

To demonstrate the performance improvements achieved by the ProtLoc-GRPO framework, we conducted extensive experiments comparing protein subcellular localization predictions using graph-based models constructed from both unpruned and ProtLoc-GRPO-pruned PPI edges. We thoroughly investigated different types of graph-based models, including GCN, GAT, and GraphSAGE, in the reward function within the ProtLoc-GRPO framework to understand how the choice of architecture influences pruning effectiveness and downstream prediction. Importantly, the architecture of the graph-based model used for final prediction is identical to the graph-based model in the reward function employed in the ProtLoc-GRPO framework. To ensure a fair and robust comparison, both the models built on unpruned and pruned PPIs were trained and evaluated using the same five-fold cross-validation splits. The difference between them lies in the PPI edges retained for graph construction, with ProtLoc-GRPO selectively removing less informative edges based on its learned pruning policy. We assessed model performance using several evaluation metrics, including macro-F1 score, macro-precision, macro-recall, and mean AUC, across five distinct human cell lines and nine subcellular localization classes. Furthermore, we systematically explored a wide range of pruning rates, varying from 10% to 80%, to examine the trade-off between graph pruning rate and prediction accuracy.

Table 1 shows that graph-based models constructed using ProtLoc-GRPO pruned PPI networks consistently outperform those built on unpruned PPI networks in protein subcellular localization prediction for all models. The best macro-F1 score is achieved by the GCN model trained on the PPI network pruned by 20%, yielding a 7% improvement compared to the GCN model trained on the unpruned PPI network. As the pruning rate increases more than 20%, the macro-F1 generally tends to decline across most metrics for all models. The GraphSAGE model built on ProtLoc-GRPO with 10% pruned PPI outperforms their unpruned counterparts in terms of mean AUC. However, ProtLoc-GRPO with different pruning rates does not always outperform its unpruned counterparts. Once the pruning rate drops below 60 percent, the prediction performance begins to decline, falling behind the models trained on unpruned PPI networks. This observation suggests that approximately 10%-20% of the PPI edges in the original network may be irrelevant or even detrimental to accurate subcellular localization prediction.

To gain a deeper understanding of ProtLoc-GRPO under the 20% pruning rate, we analyzed the mean reward score during reinforcement learning training. As shown in Figure 3, the mean reward scores for all models increase in the early stages of training, indicating that the GAT policy is being effectively optimized by the reward function. After reaching the peak reward score, all models show a decline, likely due to ProtLoc-GRPO exploring alternative PPI edges with potentially high importance. Notably, the reward scores for GraphSAGE exhibit greater fluctuation compared to the relatively stable trends observed for GAT and GCN. This suggests that ProtLoc-GRPO optimization is more stable when applied to transductive models than to inductive models. The performance of GCN built on ProtLoc-GRPO pruned PPI network at 20% pruning (detailed in the Supplementary Figure 2) shows consistent results across all five folds, with similar values for AUC, Macro-F1, macro-precision, and macro-recall between the training, validation, and test sets, indicating robust model generalization without significant overfitting or underfitting.

**Figure 3.**
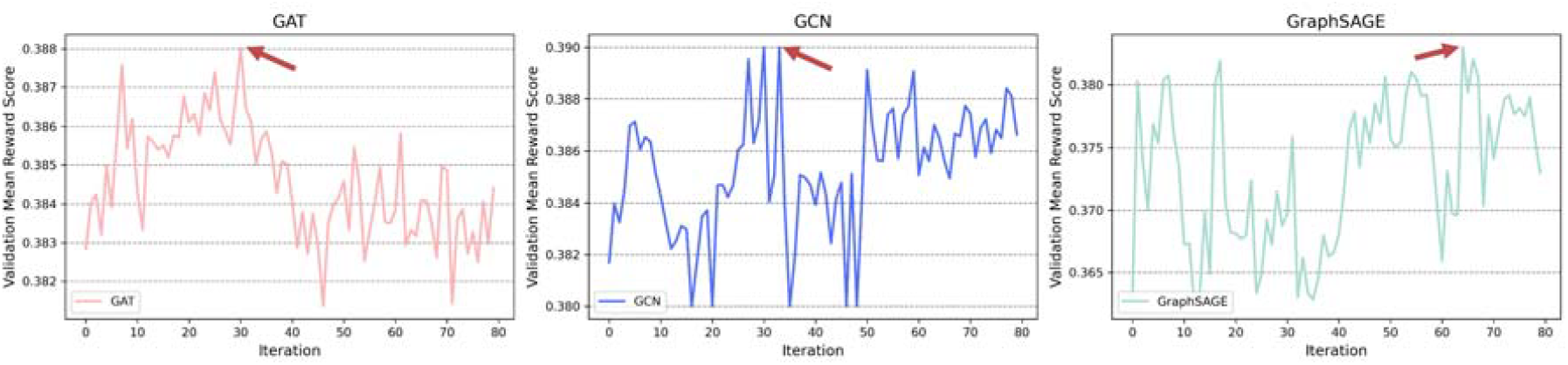
The mean reward score curve for the graph-based models built on ProtLoc-GRPO pruned PPI networks with a 20% pruning rate. The red arrow indicates the best performance on the validation set, and the policy model on those iterations will be used for evaluation.

### Pruned PPI edges compared to other important methods

In this study, we compared ProtLoc-GRPO with other edge importance estimation methods, including graph interpretation techniques and PPI confidence-based (as shown in Figure 4a). For interpretation-based approaches, we used both an attention-based method and a model-agnostic method. To obtain interpretation results, we first train a GAT model using the unpruned PPI network on the training set. In the attention-based method, we extracted attention scores^42^ from a single attention head to rank the importance of each edge using the test set. For the model-agnostic method, we applied GNNExplainer^43^ on the well-trained GAT model to estimate edge importance using the test set. The PPI confidence score-based approach relies on the scores provided by the original PPI databases. However, because these confidence scores vary in scale across different databases (as detailed in Supplementary Table 2), directly ranking edges in our globally unified PPI network becomes challenging. To address this, we designed a greedy rule-based pruning strategy. Specifically, we treated PPIs that exist in only one source as candidate edges and grouped them into four separate sets based on their source. We then pruned low-confidence interactions from each group accordingly. To compare performance, we built GAT, GCN, and GraphSAGE models using PPI networks pruned by different methods and evaluated them using a five-fold cross-validation setup.

**Table 2.** Ablation study results for the protLoc-GRPO framework. The results present the performance of the ProtLoc-GRPO framework across various combinations of its key hyperparameters: the presence or absence of KL divergence regularization, the group size utilized by the policy model, and the gradient clipping range. All experiments were conducted with a fixed pruning ratio of 20% on PPI edges, and the GCN model was consistently employed as the graph-based model during both the reward computation and evaluation stages.

| Beta | Size of group | Clip value ($\epsilon$) | AUC | Macro-F1 | Macro-precision | Macro-recall |
| --- | --- | --- | --- | --- | --- | --- |
| 0.0<br>(w/o KL) | 2 | 0.2 | 71.34 | 37.89 | 35.31 | 56.46 |
|  |  | 0.4 | 71.46 | 37.79 | 35.26 | 55.91 |
|  |  | 0.6 | 71.34 | 37.84 | 35.44 | 55.99 |
|  | 5 | 0.2 | 71.65 | **38.11** | **35.55** | 56.95 |
|  |  | 0.4 | 71.14 | 37.50 | 34.19 | 57.66 |
|  |  | 0.6 | 70.79 | 37.40 | 34.56 | 57.56 |
|  | 8 | 0.2 | 71.75 | 37.72 | 34.48 | 57.78 |
|  |  | 0.4 | **71.86** | 37.75 | 34.80 | 56.97 |
|  |  | 0.6 | 71.67 | 37.93 | 34.89 | 56.82 |
| 0.2<br>(w/ KL) | 2 | 0.2 | 71.64 | 38.09 | 35.60 | 56.47 |
|  |  | 0.4 | 71.73 | 37.93 | 34.99 | 56.55 |
|  |  | 0.6 | 71.78 | 37.99 | 35.44 | 57.18 |
|  | 5 | 0.2 | 70.90 | 37.16 | 32.54 | **59.70** |
|  |  | 0.4 | 71.75 | 37.72 | 34.48 | 57.78 |
|  |  | 0.6 | 71.75 | 37.72 | 34.48 | 57.78 |
|  | 8 | 0.2 | 71.75 | 37.72 | 34.48 | 57.78 |
|  |  | 0.4 | 71.75 | 37.72 | 34.48 | 57.78 |
|  |  | 0.6 | 71.55 | 37.86 | 34.56 | 57.50 |

**Figure 4.**
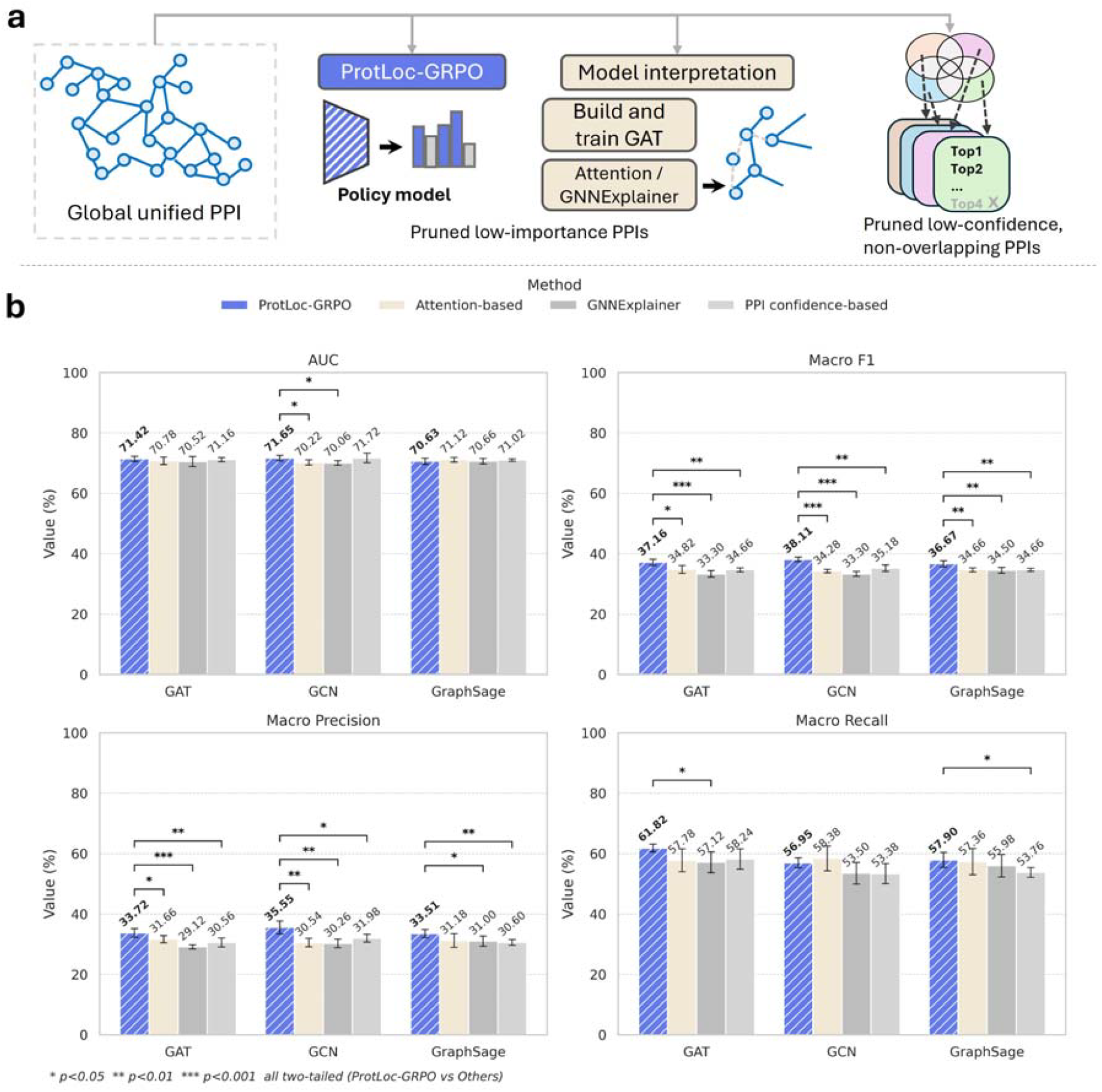
Comparison of different edge importance estimation methods. (a) Overview of different edge importance estimation methods. (b) The pruning rate is set to 20% for all methods. Each model is constructed using a PPI network pruned by its corresponding method. The bar chart presents the results of paired t-tests for each evaluation metric across different models. An asterisk (*) indicates that the performance difference between two methods is statistically significant.

Figure 4b demonstrates that the graph-based models built on ProtLoc-GRPO-pruned PPI edges outperform those constructed using other methods when the pruning rate is set to 20%. In particular, the GCN model built on ProtLoc-GRPO-pruned PPI edges shows significant improvements (*p*-value less than 0.05) in AUC, macro-F1, and macro-precision compared to the other methods. Notably, the interpretation-based approaches and the PPI confidence-based method exhibit very close and comparably lower macro-F1 performance, significantly lagging behind ProtLoc-GRPO. These results suggest that, while general explainability methods can improve model interpretability, their pruning strategies do not necessarily enhance predictive performance in protein subcellular localization tasks. In contrast, ProtLoc-GRPO directly optimizes for predictive accuracy, making it more effective at preserving essential information and eliminating noise than methods that assign importance scores post hoc.

### Representation of protein sequence

To evaluate whether ProtLoc-GRPO improves the latent space of sequence representations, we analyzed the embeddings generated by GCN models. Since a single protein may localize to multiple subcellular compartments across different cell lines, we conducted a one-vs-all analysis separately for each cell line to enable clearer visualization. Specifically, we compared the sequence embeddings produced by GCN models constructed on PPI networks pruned by ProtLoc-GRPO at a 20% pruning rate with those generated using unpruned PPI networks. Supplementary Figures 3-7 shows a UMAP^44^ visualization of the protein embeddings derived from the GCN node representations. The visualization indicates that the embeddings from the GCN model trained on the ProtLoc-GRPO-pruned PPI network exhibit better separation among subcellular localization classes compared to those from the unpruned network.

To quantitatively assess this improvement, we employed the Calinski-Harabasz Index (CHI), which evaluates clustering quality by measuring the ratio of between-cluster to within-cluster dispersion. Each subcellular localization was treated as a distinct cluster, with one localization designated as the positive class and all others grouped as the negative class. CHI scores were computed based on the sequence embeddings. Figure 5 presents the CHI scores across all cell lines. The results show that embeddings from the GCN model trained on the ProtLoc-GRPO pruned PPI network achieved higher CHI scores in 30 out of 45 subcellular localizations, highlighting the robustness of the ProtLoc-GRPO pruning strategy. The most substantial CHI improvements resulting from ProtLoc-GRPO pruning were frequently observed in major organelles such as the mitochondria, nucleus, and endoplasmic reticulum. The greatest improvement was a 33% increase in CHI for the endoplasmic reticulum in the U-251MG cell line. These findings indicate that ProtLoc-GRPO enhances both the quality and discriminative power of protein sequence embedding.

**Figure 5.**
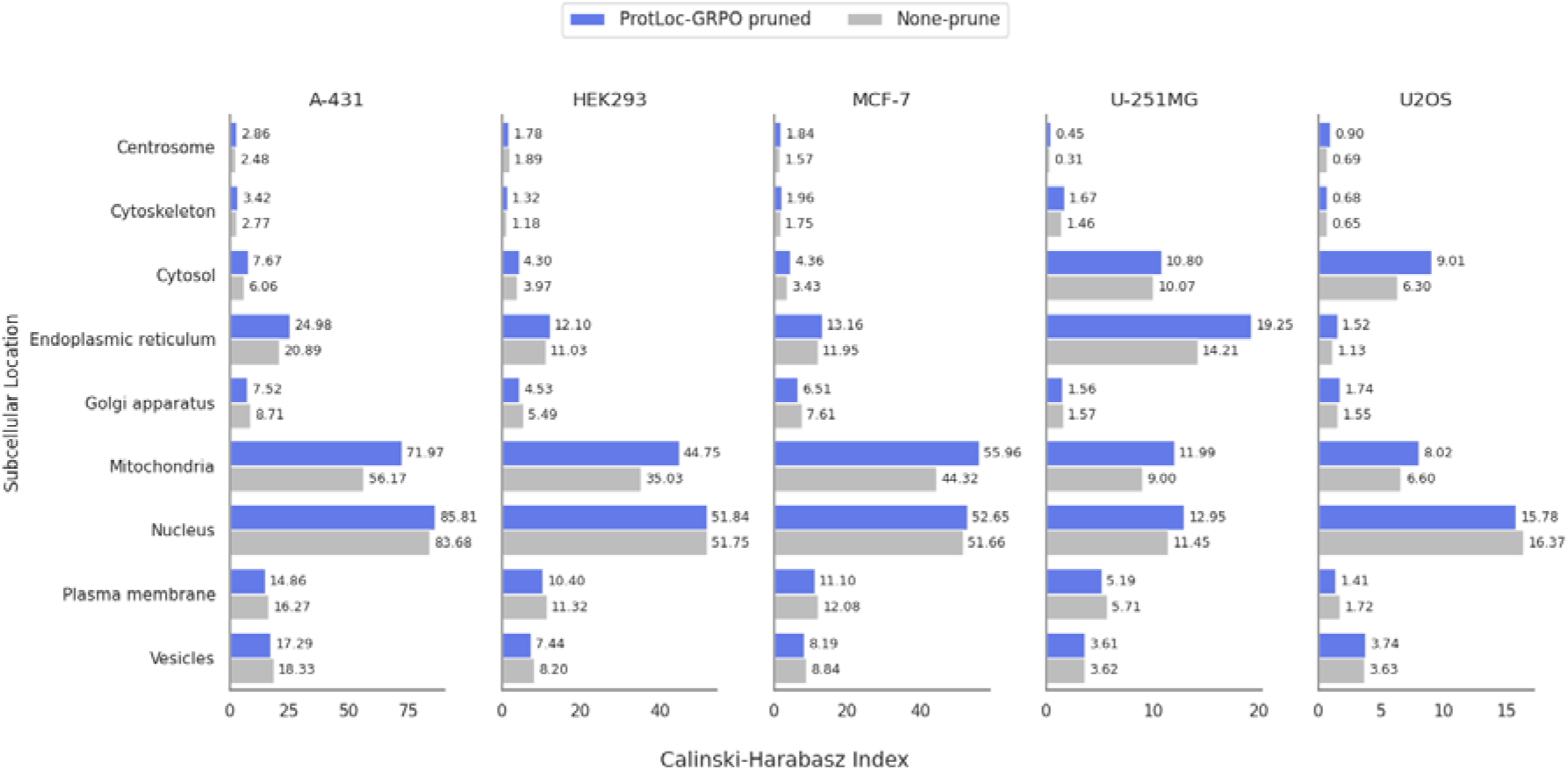
Calinski-Harabasz index of one-vs-others protein clusters for subcellular localization across different cell lines. This figure compares the CHI for protein clusters representing specific subcellular locations against all other locations within five distinct cell lines. Each subplot shows the CHI for proteins associated with various subcellular localizations, contrasting performance with ProtLoc-GRPO pruned PPI networks (blue bars) versus unpruned PPI networks (gray bars). A higher CHI indicates more compact and well-separated protein clusters, demonstrating that ProtLoc-GRPO pruning generally improves the discriminative quality of protein groupings by subcellular localization.

### Ablation Study

We performed ablation studies to investigate the contribution of key components in the ProtLoc-GRPO framework, including the group size used by the policy model, the presence or absence of KL divergence regularization, and the gradient clipping range. For all ablation experiments, the pruning ratio of PPI edges was fixed at 20%, and the graph-based model in the reward function and test stage is GCN.

Table 2 reports the performance of various combinations of the three key components within the ProtLoc-GRPO framework. The configuration incorporating KL divergence regularization demonstrates consistently stable performance, as the KL term constrains policy updates by penalizing deviations from the previous policy, thereby limiting excessive exploration. In contrast to the typical post-training setting of GRPO in LLM, ProtLoc-GRPO exhibits improved performance as the group size decreases. The improvement is attributed to the use of the macro-F1 score as a reward signal, which offers continuous and informative feedback to the policy model, thereby mitigating the risk of local minima that commonly arise with a sparse reward signal^45,46^. Additionally, the clipping parameter plays a crucial role in regulating the degree of exploration during training. Larger clipping values allow for more aggressive policy updates, which may lead to instability in the reinforcement learning process.

## Discussion

In this study, we present ProtLoc-GRPO, a novel reinforcement learning framework designed to improve cell line-specific protein subcellular localization prediction by optimizing the structure of the underlying PPI network. Our approach addresses a critical limitation of current PPI networks, namely the presence of systematic noise and biologically irrelevant interactions, which can hinder prediction accuracy. Leveraging the GRPO framework, ProtLoc-GRPO learns to identify and retain only the most informative and predictive PPI edges, effectively filtering out misleading or non-contributory connections. To our knowledge, this work is the first to predict cell line-specific protein subcellular localization and the first to apply the GRPO framework to graph-based models in bioinformatics tasks. Unlike most GRPO applications that optimize LLM policies to enhance language model capabilities for specific tasks, we optimize a GAT policy to score PPI edges and prune those with the lowest importance.

Our empirical results demonstrate the efficacy of ProtLoc-GRPO. Graph-based models utilizing ProtLoc-GRPO pruned PPI networks consistently outperform those based on unpruned networks. An optimal pruning rate of 20% was identified, yielding a notable 7% improvement in macro-F1 for the GCN model. Furthermore, ProtLoc-GRPO outperformed other pruning strategies, including attention-based, GNNExplainer, and PPI confidence-based methods. At the 20% pruning rate, ProtLoc-GRPO consistently achieved higher AUC, macro-F1, macro-precision, and macro-recall. Interestingly, interpretability-based pruning methods did not translate into superior predictive performance, often yielding results comparable to those of confidence-based pruning. This underscores that direct optimization for prediction is more effective than relying on general importance metrics for network refinement. The improved network structure also enhanced latent space representations. The Calinski-Harabasz Index for one-vs-others clustering demonstrated that ProtLoc-GRPO pruning consistently resulted in better-defined and more separable protein clusters across diverse subcellular locations and cell lines. Ablation studies confirmed that without the KL divergence regularization, it helps achieve higher peak performance in terms of macro-F1. Unlike GRPO applications in LLMs, ProtLoc-GRPO remains effective even with smaller group sizes, owing to its use of a continuous macro-F1 reward signal. However, the limitation of the method is its reliance on the initial PPI network, as proteins not represented in the original interaction graph cannot benefit from the optimization process. Another limitation is that the pruning rate is not optimized by ProtLoc-GRPO, which suggests that the method could be redesigned to automatically learn or adaptively optimize the pruning rate.

While our work in this study focuses on improving protein localization predictions, the challenges we address, particularly the identification of high-value interactions from noisy biological networks, are broadly relevant across a wide range of bioinformatics applications. Beyond subcellular localization, our approach has the potential to enhance data quality and model robustness in various contexts. For example, protein function annotation often relies on interaction partners with known functions^47^, and disease gene prioritization depends heavily on identifying genes that are functionally related to known disease-associated genes within a network^48^. An optimized interaction network provides a more accurate biological context for inferring such relationships, thereby improving the reliability of downstream predictions.

## Methods

### Dataset

#### Protein-protein interaction data

Our unified PPI network is constructed as the union of physically validated interactions from BioGRID, the Human Reference Interactome (HuRI), STRING (for human), and Menche et al., resulting in a total of 15,461 nodes and 407,641 edges. Each source employs distinct methodologies for curating and validating physical protein–protein interactions. BioGRID, HuRI, and Menche et al. are based on well-established and widely cited datasets of experimentally validated human PPIs. STRING integrates PPIs from a variety of sources, including experiments, curated databases, text mining, and co-expression analysis. Because STRING contains a large number of low-confidence interactions that can overshadow other datasets, we removed edges with a confidence score lower than 0.7. By integrating these four datasets and filtering accordingly, we constructed a globally unified and high-confidence PPI network for our study.

#### Cell line-specific protein subcellular localization dataset

Our study utilizes subcellular localization data provided by version 23 of the Human Protein Atlas (HPA)^1^. Protein sequences were obtained from either the UniProt^49,50^ or Ensembl^51^ database. In total, 13,534 genes with subcellular localization information were initially collected from 41 cell lines. Due to the limited number of proteins annotated in most cell lines, we restricted our analysis to proteins with subcellular information in cell lines containing more than 1,000 annotated proteins. This filtering resulted in 11,244 proteins across 5 cell lines. We further excluded proteins not presenting in the global PPI network, yielding a final set of 9,713 proteins. For localization labels, we consolidated the official 33 subcellular localizations into 9 broader categories to address the limitations of the training dataset (as detailed in Supplementary Table 3).

#### Evaluation and metrics

We divided our collected dataset into five folds using ggsearch^52^, ensuring that the sequence identity between proteins in different folds remained below 30%. During the five-fold cross-validation, three folds were used for training the policy, a single fold was used for reward calculation, and the remaining fold was held out for final testing. For each cell line and subcellular localization category, we calculated the AUC score at each position and reported the mean AUC. Additionally, we evaluated model performance using macro-F1, macro-precision, and macro-recall metrics across all subcellular localizations and cell lines.

Subcellular locations with no labeled proteins were excluded from the evaluation metrics to ensure reliable performance assessment.

### ProtLoc-GRPO framework

#### GAT policy model

Similar to the GRPO framework, we design a GAT policy model. The policy model π_θ_ estimates the importance *s*_*i*_ of each edge *e*_*i*_ in the PPI network *G* using the GAT model *f*_θ_, which takes node embeddings of proteins 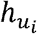 and 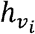 as input.

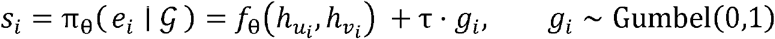

During the training stage, a Gumbel noise value *g*_*i*_, sampled from the standard Gumbel distribution, is added to the corresponding importance score *s*_*i*_ for each edge, similar to previous works using Gumbel-based sampling^53,54^. The temperature parameter τ scales the noise magnitude to control the balance between exploration and exploitation during sampling. We fixed τ to 0.001. The process of adding Gumbel noises is conducted *G* times to sample *G* different sets of PPI edges. In each sampled set, the top-*k* percent of edges with the highest importance scores are retained, while the remaining edges are pruned. The policy model outputs *G* subgraphs of the PPI network, each corresponding to a distinct pruning instance. During inference, the top-*k* percent of edges with the highest importance scores are selected deterministically by setting τto 0, without adding Gumbel noise as done during training. As a result, a single pruned subgraph of the PPI network is generated.

#### Reward function

The goal of ProtLoc-GRPO is to prune less important PPI edges and improve protein subcellular localization prediction. Unlike the reward function used in most LLM post-training^33,41,55,56^, which are either rule-based or rely on pretrained scoring functions, our reward function in ProtLoc-GRPO is similar to a general machine learning framework that includes both training and evaluation stages. Specifically, the reward function includes a graph-based model constructed using a subgraph of the PPI network generated by the policy model. This graph-based model is trained using a binary cross-entropy loss on the training dataset. It is then evaluated on a validation dataset, and the macro-F1 score is used as the final reward. For the *G* different subgraphs {*E*_*1*_, *E*_*2*_, …, *E*_*G*_} produced by the policy model, we train and evaluate *G* separate graph-based models using the same data. The resulting macro-F1 scores are normalized by *z*-score and used to update the policy model through the GRPO objective function. As the KL divergence regularization was removed, the reference model was excluded from the training process.

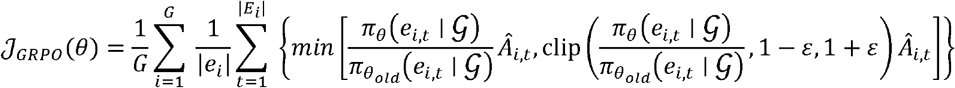

Where _*ei,t*_, represents the *t*-th edge in the *i*-th PPI network *G*, *Â*_*I,t*_,(the advantage) is calculated based solely on the relative rewards of the outputs within each group, and ε is a hyperparameter.

### Graph Network

#### Node features

We used the ESM2-650M^57^ protein language model to extract high-dimensional embeddings as node features for the graph-based policy model and reward function. The ESM2-650M model takes a protein sequence as input. We truncated all protein sequences to 1022 amino acids due to the ESM2 model pretrained on the protein sequence of up to 1024 tokens (including two special tokens CLS and EOS). The sequences shorter than 1022 amino acids are padded to align with the length of the input. We remove the special tokens (CLS and SEP) introduced by the ESM2-650M, as well as any padding tokens, and retain a sequence of hidden states *h* with the same length as the original input sequence *s*.

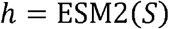

To ensure that the graph representation has a consistent input size regardless of the protein sequence length, we then compute the mean over the hidden states.

### Graph-based models

We conduct comprehensive experiments by integrating GAT, GCN, and GraphSAGE into the reward function of the ProtLoc-GRPO framework. For the policy model, we adopt GAT architecture followed by fully connected layers to estimate the importance of PPI edges, leveraging its attention mechanism to capture the relative significance of neighboring nodes. For the reward function, we evaluate GAT, GCN, and GraphSAGE, as pruning edges may affect each graph model differently in terms of localization prediction performance, necessitating a thorough comparison. These models are also followed by fully connected layers to perform protein subcellular localization prediction. The output consists of 45 probabilities per sample, corresponding to 9 localization categories across 5 distinct cell lines. To convert the predicted probabilities into binary labels, we determine an optimal threshold for each of the 45 outputs by maximizing the F1 score on the validation set.

## Author Contributions

S.Z., W.Z., and C.L. contributed to the development of the framework and drafting of the manuscript. D.X., Y.J., and S.Z. conceived the study. Y.J., D.W., and Q.S. participated in discussions throughout the project. All authors revised the manuscript. D.X. provided overall guidance to this study.

## Acknowledgements

This study was funded by the National Institutes of Health (grants R35GM126985 and R01LM014510). This work used the GPU resources through allocations NAIRR240246 and NAIRR250134 from the National Artificial Intelligence Research Resource (NAIRR).

## Data Availability

All protein sequence data used in this study are available from UniProt and Ensembl. Cell line-specific subcellular localization annotations can be obtained from the Human Protein Atlas (HPA).

## Code Availability

The Python code developed for this work, along with documentation and usage examples, is publicly available on GitHub at https://github.com/shuaizengMU/ProtLoc-GRPO.

