## Supplementary for "ProtLoc-GRPO: Cell line-specific subcellular localization prediction using a graph-based model and reinforcement learning"

**Supplementary Table 1**. Overview of selected cell lines and corresponding tissue of origin.

| Cell Line | Tissue |
|---|---|
| A-431 | Skin |
| U2OS | Bone |
| U-251MG | Brain |
| MCF-7 | Pleural effusion |
| HEK293 | Kidney |

**Supplementary Table 2**. Characteristics of PPI datasets with a rule applied for identifying and filtering out low-confidence PPIs. The NA represents no confidence scores in the corresponding PPI dataset.

| PPI dataset | Number of edges | Range/label of confidence | Rule for low confidence PPI |
|---|---|---|---|
| *BioGRID* | 13,2767 | NA, 0~1000 | PPI with confidence value (NA) |
| *HuRI* | 25,558 | NA | PPI with confidence value (NA) |
| *Menche et al.* | 98,021 | binary, complexes, metabolic, signaling, literature | binary, complexes, metabolic, signaling |
| *STRING* | 97,577 | 0.7~1.0 | Lower value represents low confidence |

**Supplementary Table 3**. Mapping of HPA subcellular localization annotations to merged categories.

| # | Our merged subcellular | HPA subcellular |
|---|---|---|
| 1 | Nucleus | Nuclear membrane |
| | | Nucleoli |
| | | Nucleoplasm |
| 2 | Cytoskeleton | Cytosol |
| | | Actin filaments |
| | | Intermediate filaments |
| 3 | Endoplasmic reticulum | Endoplasmic reticulum |
| 4 | Golgi apparatus | Golgi apparatus |
| 5 | Plasma membrane | Plasma membrane |
| 6 | Vesicles | Vesicles |
| 7 | Centrosome | Centrosome |
| 8 | Microtubules | Microtubules |
| 9 | Mitochondria | Mitochondria |

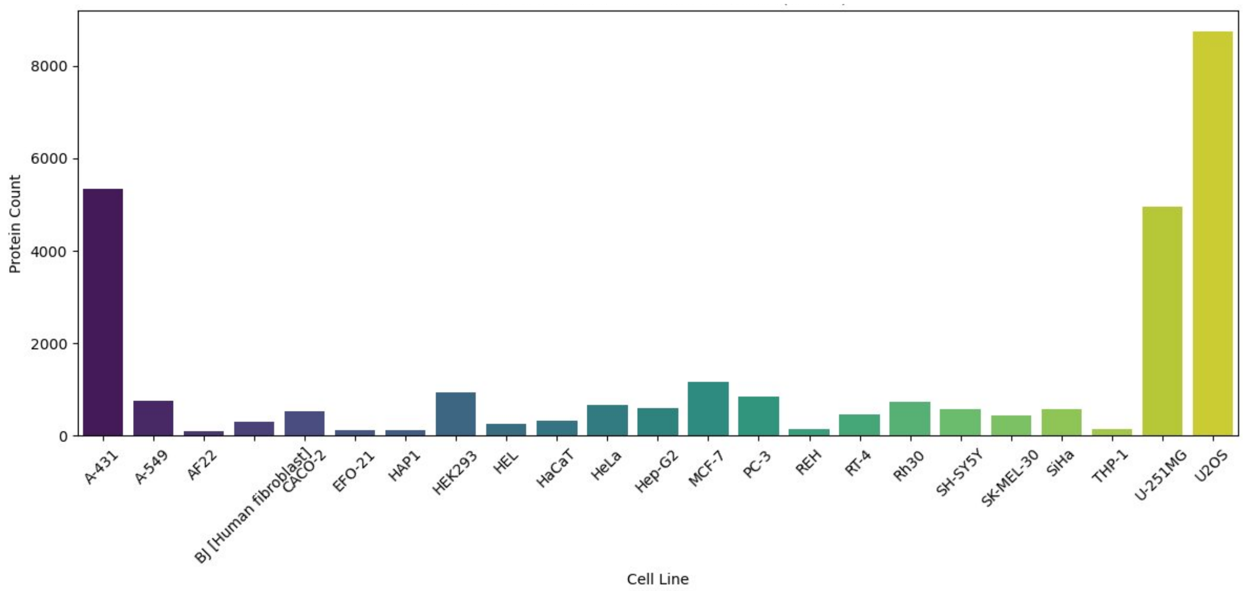


**Supplementary Figure 1**. Distribution of protein counts across cell lines from HPA.

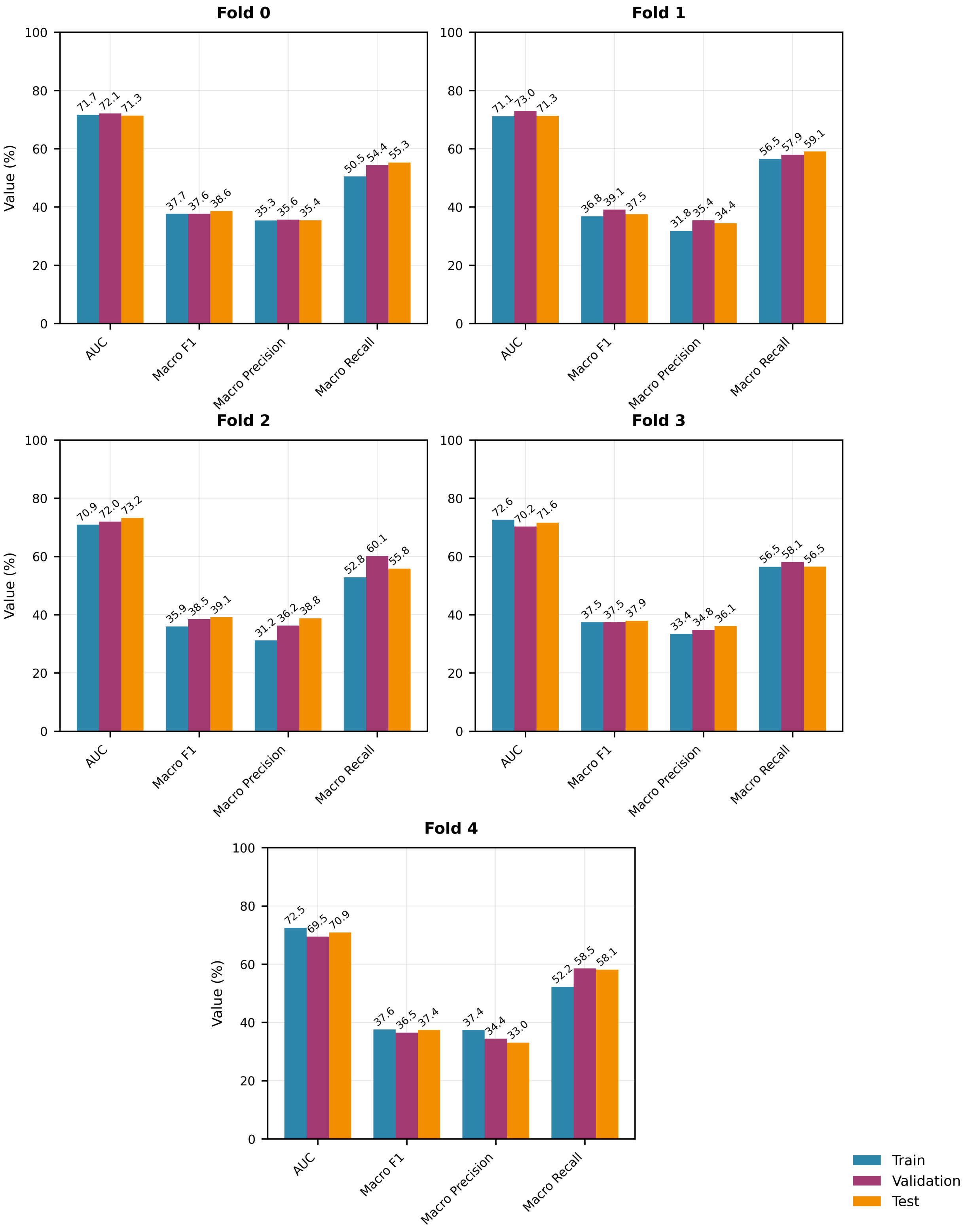


**Supplementary Figure 2**. Performance on training, validation, and test datasets across all five folds.

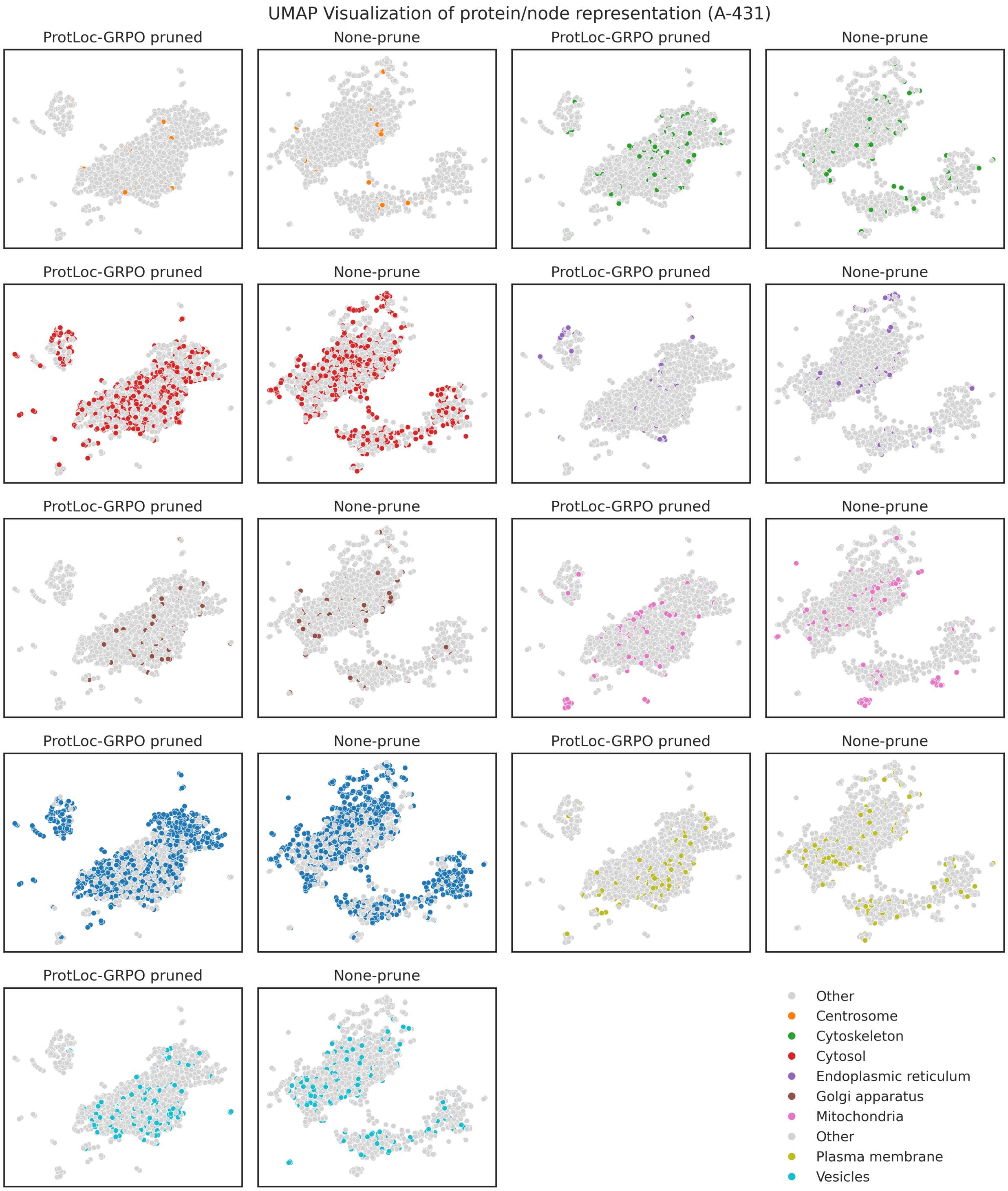


**Supplementary Figure 3**. UMAP visualization of protein representation in A-431 cell line. Each subplot displays the protein representations from either ProtLoc-GRPO pruned PPI networks or non-pruned PPI networks. Each data point represents a protein embedding. In each subplot, colored points highlight a particular subcellular location, while gray points represent all other locations.

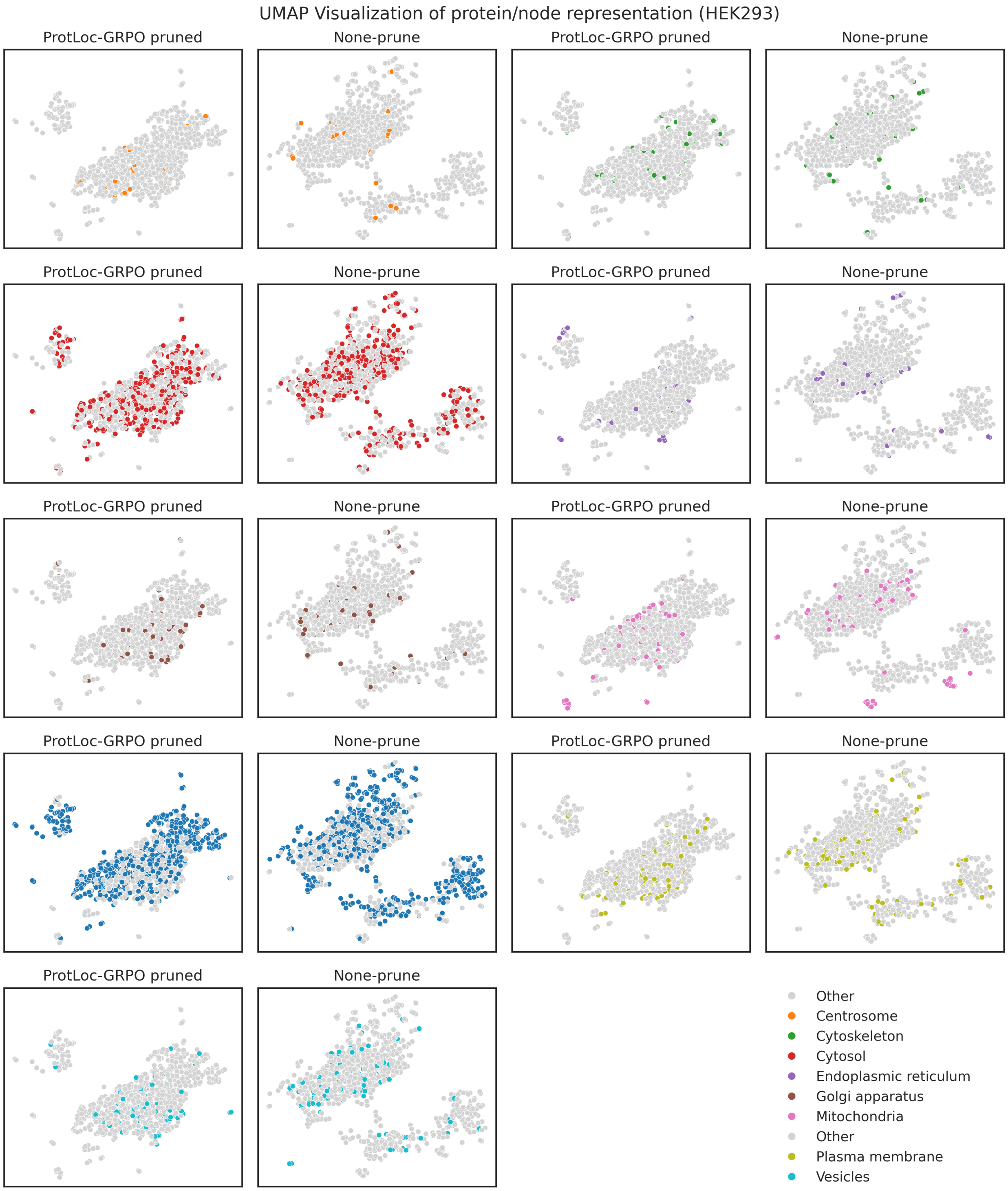


**Supplementary Figure 4**. UMAP visualization of protein representation in the HEK293 cell line. Each subplot displays the protein representations from either ProtLoc-GRPO pruned PPI networks or non-pruned PPI networks. Each data point represents a protein embedding. In each subplot, colored points highlight a particular subcellular location, while gray points represent all other locations

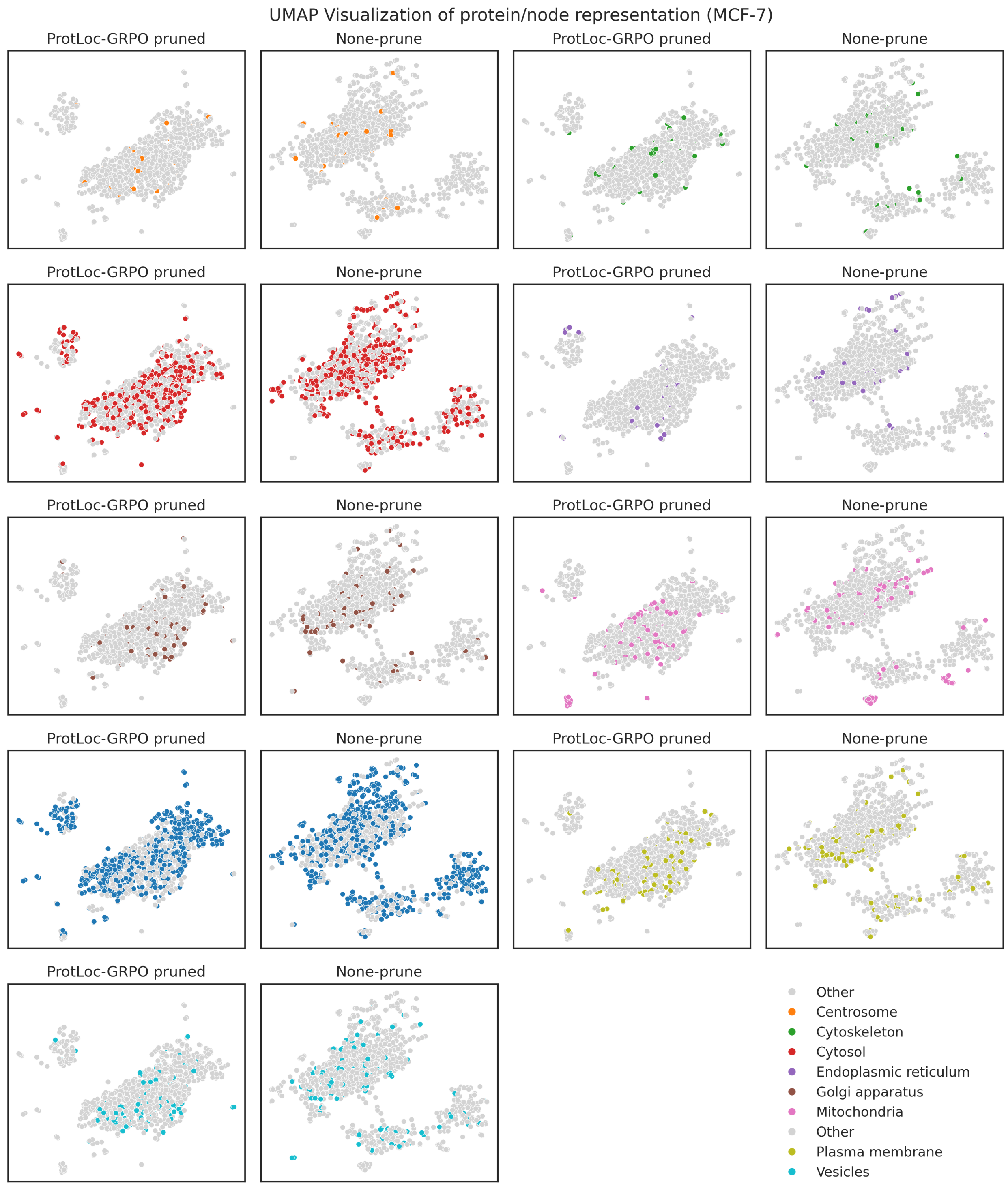


**Supplementary Figure 5**. UMAP visualization of protein representation in the MCF-7 cell line. Each subplot displays the protein representations from either ProtLoc-GRPO pruned PPI networks or non-pruned PPI networks. Each data point represents a protein embedding. In each subplot, colored points highlight a particular subcellular location, while gray points represent all other locations

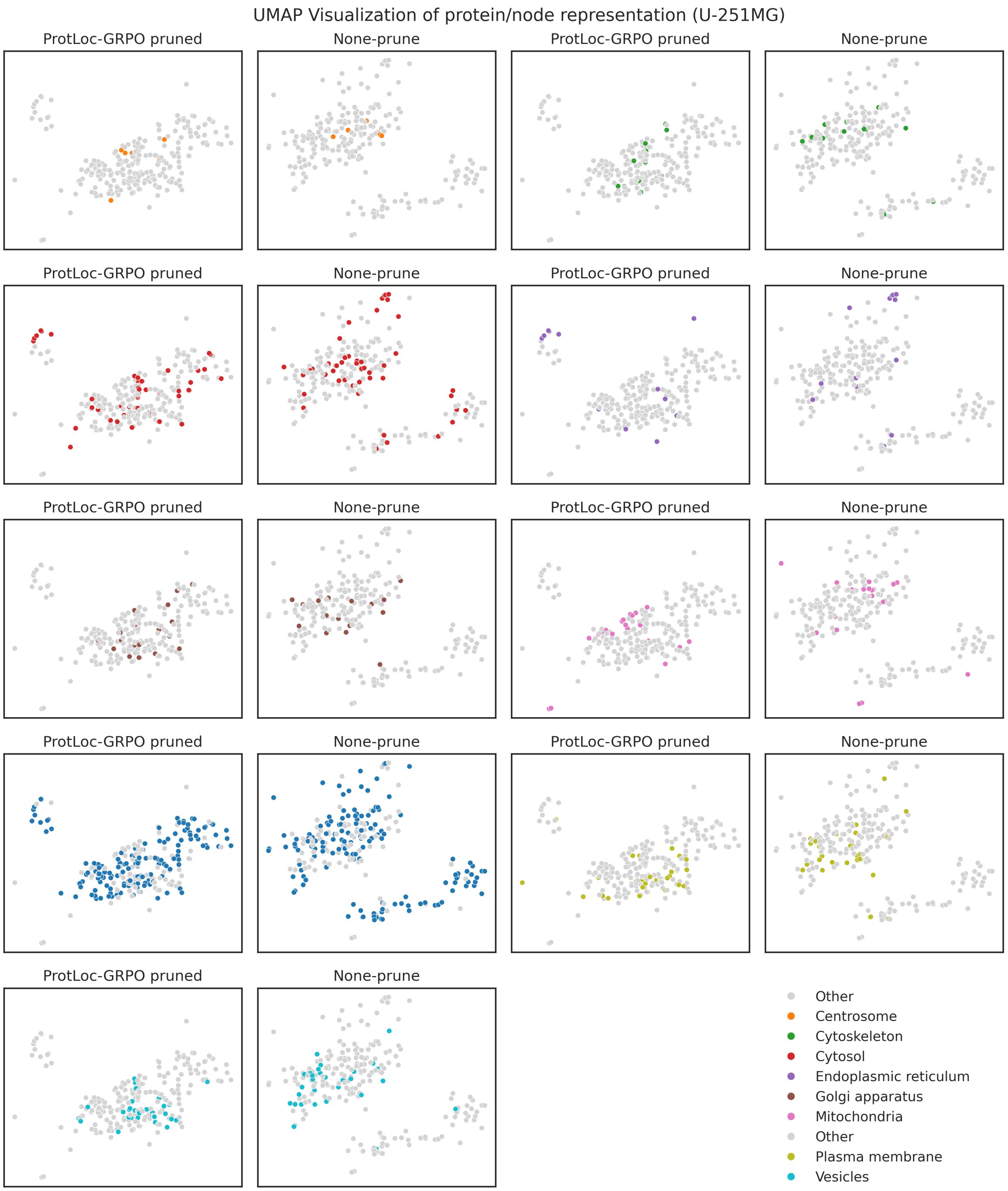


**Supplementary Figure 6**. UMAP visualization of protein representation in the U-251MG cell line. Each subplot displays the protein representations from either ProtLoc-GRPO pruned PPI networks or non-pruned PPI networks. Each data point represents a protein embedding. In each subplot, colored points highlight a particular subcellular location, while gray points represent all other locations

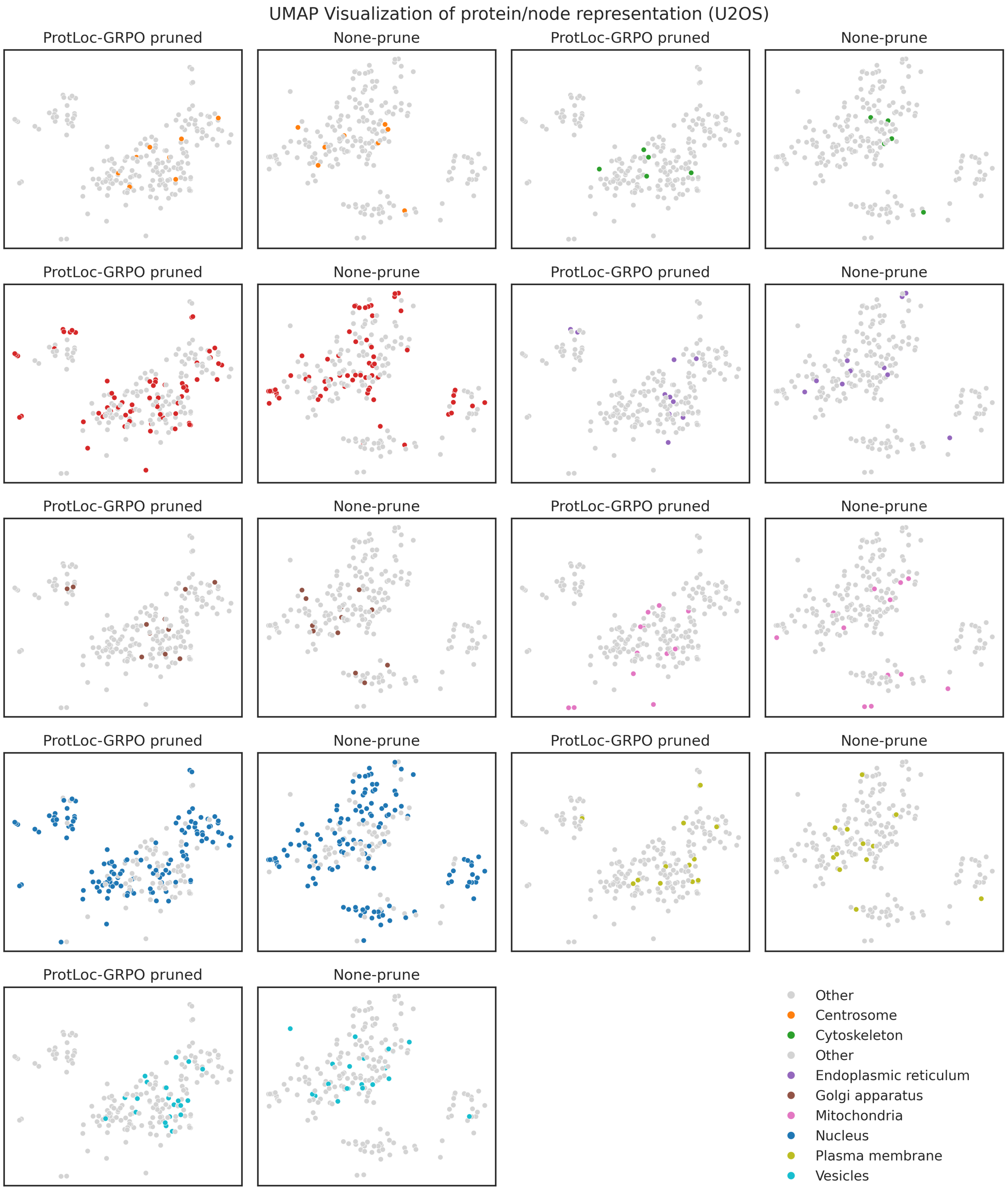


**Supplementary Figure 7**. UMAP visualization of protein representation in the U2OS cell line. Each subplot displays the protein representations from either ProtLoc-GRPO pruned PPI networks or non-pruned PPI networks. Each data point represents a protein embedding. In each subplot, colored points highlight a particular subcellular location, while gray points represent all other locations
